# In Vitro Inhibition of SARS-CoV-2 Infection by Bovine Lactoferrin

**DOI:** 10.1101/2020.05.13.093781

**Authors:** Carlos Alberto Marques de Carvalho, Aline da Rocha Matos, Braulia Costa Caetano, Ivanildo Pedro de Sousa Junior, Samir Pereira da Costa Campos, Barbara Rodrigues Geraldino, Caroline Augusto Barros, Matheus Augusto Patricio de Almeida, Vanessa Pimenta Rocha, Andréa Marques Vieira da Silva, Juliana Gil Melgaço, Patrícia Cristina da Costa Neves, Tamiris Azamor da Costa Barros, Ana Paula Dinis Ano Bom, Marilda Mendonça Siqueira, Sotiris Missailidis, Rafael Braga Gonçalves

## Abstract

Since its emergence in late 2019, severe acute respiratory syndrome coronavirus 2 (SARS-CoV-2) has been posing a serious threat to public health worldwide as the causative agent of coronavirus disease 2019 (COVID-19). Now distributed in a pandemic pattern, this disease still lacks an effective drug treatment with low toxicity, leading pharmaceutical companies and research labs to work against time to find a candidate molecule to efficiently treat the affected patients. Due to the well-known broad-spectrum antimicrobial activity of the lactoferrin protein, we sought to verify whether its bovine form (bLf) would also be effective in vitro against SARS-CoV-2. Using an antiviral assay based on quantitative reverse transcription-polymerase chain reaction (qRT-PCR), we found that bLf reduced progeny virus yield by up to ∼84,6% in African green monkey kidney epithelial cells (Vero E6) and ∼68,6% in adenocarcinomic human alveolar basal epithelial cells (A549) at 1 mg/mL, a concentration previously shown to have low cytotoxicity. Therefore, our preliminary data suggest that bLf has the potential to constitute a biochemical approach to fight the new coronavirus pandemic.

## Introduction

Coronaviruses (CoVs) are enveloped viruses with a positive-sense, single-stranded ribonucleic acid (RNA) genome, structurally recognized by electron microscopy for their distinctive crown-shaped spike projections (Li, 2016). Although some CoVs (e.g., HKU1, NL63, 229E and OC43) were already known to cause respiratory infections in humans, this group of viruses was not considered to be highly pathogenic until the outbreaks of severe acute repiratory syndrome (SARS) in 2002 and Middle East respiratory syndrome (MERS) in 2012 (Cui et al., 2019). The so-called SARS-CoV and MERS-CoV were each responsible for hundreds of human deaths (Su et al., 2016).

In late 2019, a new CoV emerged in China and quickly spread across the world causing a respiratory illness named coronavirus disease 2019 (COVID-19) (Jiang et al., 2020). This virus, called SARS-CoV-2 due to its close phylogenetic relationship with the CoV from the 2002 outbreak, has reached a pandemic distribution pattern and has already caused thousands of human deaths worldwide (Wu et al., 2020).

Despite being less lethal than the CoVs associated to SARS and MERS, the CoV responsible for COVID-19 is more contagious than those viruses (Meo et al., 2020). The high rate of hospitalizations resulting from this new disease has caused the collapse of health systems in several countries, raising the need for an effective drug with low toxicity to treat human infections (Velavan & Meyer, 2020). In this sense, lactoferrin (Lf) – an iron-binding protein naturally found in milk and other bodily secretions with broad-spectrum antimicrobial activity (Jenssen & Hancock, 2009) – may represent an alternative.

Lf exhibits inhibitory activities against a wide range of viruses in vitro, including those responsible for respiratory tract infections such as common cold and influenza (Wakabayashi et al., 2014). In its bovine form (bLf), the protein was also shown to be effective in vitro against a pseudotyped version of the etiological agent of SARS, interfering with virus infection at the attachment stage (Lang et al., 2011).

In view of the above, this brief work aimed to assess the activity of bLf against SARS-CoV-2 in vitro, using African green monkey kidney epithelial (Vero E6) and adenocarcinomic human alveolar basal epithelial (A549) cell lines.

## Material and Methods

Vero E6 and A549 cells were cultured in Dulbecco’s modified Eagle medium (DMEM) supplemented with 10% fetal bovine serum (FBS) plus penicillin/streptomycin and incubated at 37 °C in a 5% CO_2_ atmosphere. SARS-CoV-2 was isolated after two passages of a nasopharyngeal swab sample from a positive case in Rio de Janeiro, Brazil, in Vero E6 cells. All virus-related procedures were performed in multi-user facilities with biosafety level 3 (BSL-3), according to World Health Organization (WHO) guidelines. BLf was obtained from Art’Gerecht.

For the antiviral assays, cells were plated at a density of 5 × 10^4^ cells/well in 48-well plates. After culturing overnight, cells were washed with phosphate-buffered saline (PBS) and incubated with bLf in FBS-containing DMEM at the indicated concentrations for 1 h. Then, the previous solution was removed and cells were incubated with SARS-CoV-2 at a multiplicity of infection (MOI) of 0.01 in the presence of bLf in pure DMEM for an additional 1 h. Afterwards, the virus inoculum was removed and cells were incubated with bLf-containing medium for 72 h.

Quantitative reverse transcription-polymerase chain reaction (qRT-PCR) assays were performed after extraction of viral RNA from the culture supernatant using the QIAGEN QIAamp Viral RNA Mini Kit, according to the manufacturer’s instructions. Amplification was carried out by using the QIAGEN OneStep RT-PCR Kit on the ABI PRISM 7500 System with primers and probes specific for SARS-CoV-2 as published elsewhere (Corman et al., 2020). A standard curve with a pre-quantified reference sample was used for RNA quantification, expressed as a function of volume (copies/mL).

Statistical analyses were performed using ordinary one-way ANOVA with Dunnett’s post-test for multiple comparisons on GraphPad Prism 6 Software. P-values less than 0.05 were considered as indicative of statistically significant differences.

## Results

As an initial assessment of the inhibitory activity of bLf on SARS-CoV-2 infection in a both suceptible and permissive cell line, Vero E6 cells were infected in the absence or presence of the protein at concentrations ranging from 0.2 to 1.0 mg/mL and virus load in the culture supernatant was determined by qRT-PCR. As observed, bLf promoted a dose-dependent inhibition of SARS-CoV-2 yield, which was reduced by up to ∼84,6% at 1 mg/mL based on viral RNA level (Figure 1).

**Figure 1:**
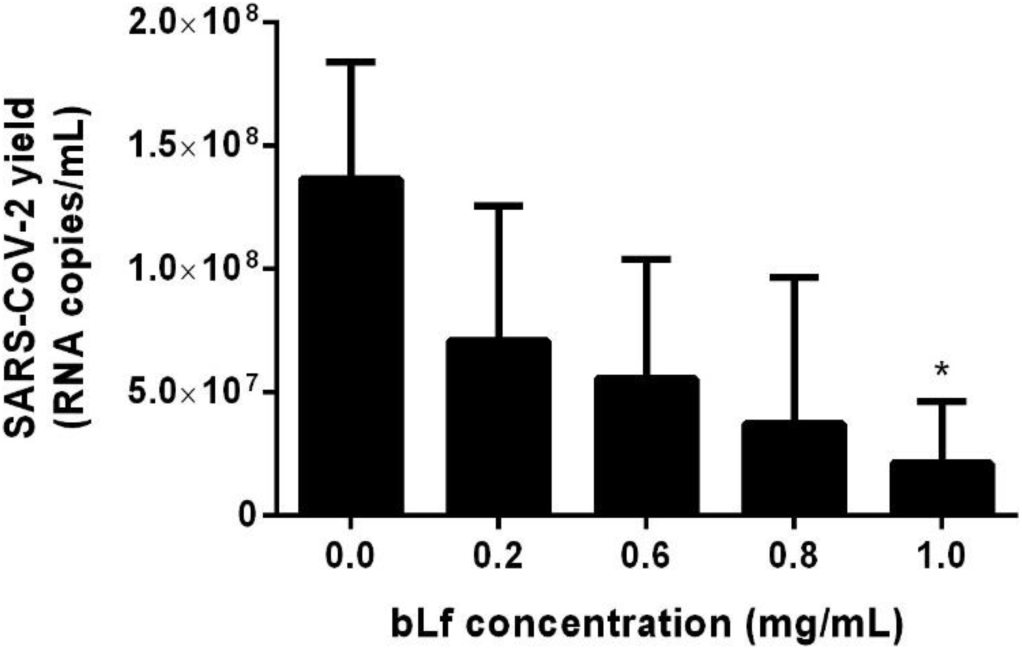
Effect of bLf on SARS-CoV-2 yield in Vero E6 cells. Cells were incubated with the indicated concentrations of bLf before, during and after virus adsorption. At 72 h post-infection, viral RNA was extracted from the culture supernatant and quantified by qRT-PCR. Data represent means plus standard deviations from 3 independent experiments and asterisk denotes a statistically significant difference in relation to control.

Next, to check if such antiviral action of bLf is also exerted on SARS-CoV-2 infection in a cell line more related to the main target organs of the virus in humans (i.e., the lungs), the previous in vitro approach based on qRT-PCR was conducted with A549 cells. Once again, bLf promoted a significant inhibitory effect on SARS-CoV-2 yield, reducing viral RNA level in the culture supernatant by up to ∼68,6% at 1 mg/mL (Figure 2).

**Figure 2:**
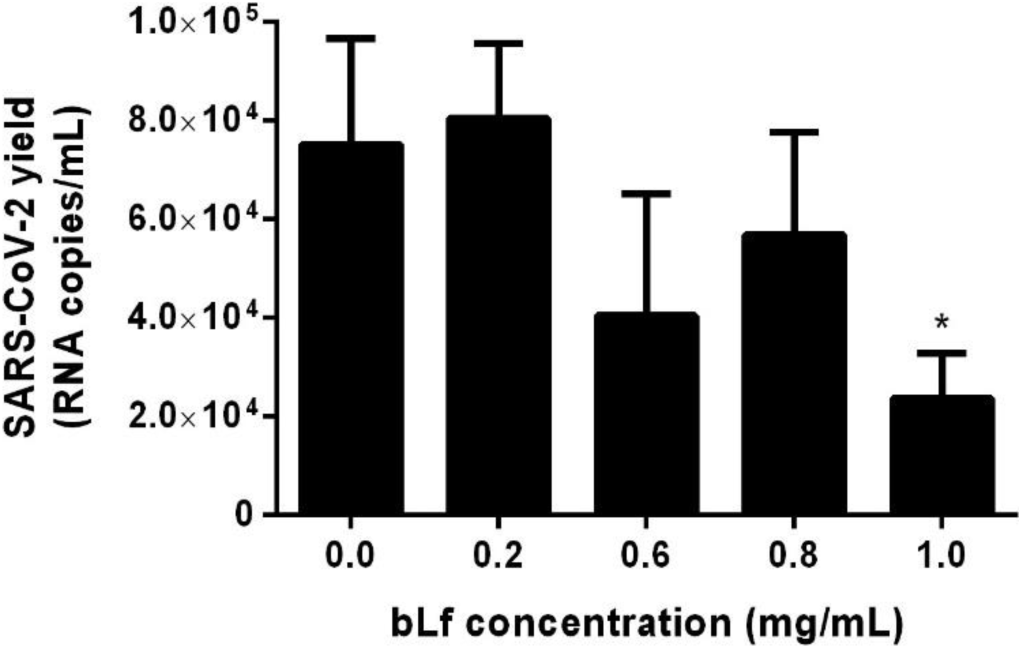
Effect of bLf on SARS-CoV-2 yield in A549 cells. Cells were incubated with the indicated concentrations of bLf before, during and after virus adsorption. At 72 h post-infection, viral RNA was extracted from the culture supernatant and quantified by qRT-PCR. Data represent means plus standard deviations from 3 independent experiments and asterisk denotes a statistically significant difference in relation to control.

## Discussion

Supportive care remains the most important management strategy for human infection by highly pathogenic CoVs, as there is currently no specific antiviral treatment that has been proven to be effective in randomized controlled trials (Zumla et al., 2016). Facing the rampant spread of COVID-19 around the world, this study demonstrated that bLf is able to significantly interfere with SARS-CoV-2 replication in vitro at 1 mg/mL. Previous studies using Vero E6 (Murphy et al., 2000) or A549 (Tung et al., 2013) cells have reported low cytotoxic effects for bLf at this concentration.

In line with a recent work on SARS-CoV-2 (Harcourt et al., 2020), our qRT-PCR data revealed that the virus replicates at higher titers in Vero E6 cells than in A549 cells, which differed by 4 log_10_. This variation may be due to a limited expression of viral subgenomic RNAs during CoV replication in the latter cells (Gillim-Ross et al., 2004).

Based on a previous work addressing the mechanism involved in the activity of bLf against a SARS pseudovirus (Lang et al., 2011), it is likely that inhibition of SARS-CoV-2 infection occurs at the entry level by blocking heparan sulfate proteoglycans (HSPGs), which are commonly exploited by viruses for cell attachment (Cagno et al., 2019). Indeed, this seems to be a general mechanism underlying the inhibitory effect of bLf on several unrelated viruses, including respiratory viruses (Superti et al., 2019) and even endemic (Carvalho et al., 2014) or epidemic (Carvalho et al., 2017) arboviruses.

Regarding in vivo responses to CoVs, it is worth noting that genes coding for Lf are highly up-regulated in patients with SARS and the protein may further inhibit virus infection in vivo by enhancing natural killer (NK) cell activity and stimulating neutrophil aggregation and adhesion (Reghunathan et al., 2005).

Recently, a prospective observational study with COVID-19 patients has shown that oral treatment with liposomal bLf allows a complete and fast recovery from the disease within the first 4-5 days, although it has not been proved that the protein acts directly on virus infection (Serrano et al., 2020). Despite being preliminary, our data provides a proof of concept to support further trials with bLf in the pandemic scenario of COVID-19.

## Ethical Statement

This study made no use of human or vertebrate animal subjects and/or tissues. All authors approved the current version of this compuscript and consented to its publication as a preprint on bioRxiv.

